# Modulation of Temporoammonic-CA1 Synapses by Neuropeptide Y is Through Y1 Receptors

**DOI:** 10.1101/2024.03.26.586875

**Authors:** Mariana A. Cortes, Aundrea F. Bartley, Qin Li, Taylor R. Davis, Stephen E. Cunningham, Patric J. Perez, Lynn E. Dobrunz

## Abstract

The reduction of neuropeptide Y (NPY), an abundant neuromodulator in the brain, has been implicated in multiple neuropsychiatric disorders, such as depression and post-traumatic stress disorder (PTSD). The CA1 region of hippocampus is an important area for anxiety and highly expresses NPY. Injection of NPY into the CA1 is anxiolytic and has been shown to alleviate behavioral symptoms in a model of traumatic stress. It is known that activation of NPY Y1 receptors has anxiolytic effects and that NPY’s anxiolytic effects in CA1 are blocked by an NPY Y1 receptor antagonist. However, the location of Y1 receptors mediating NPY’s anxiolytic effects in CA1 is not yet known. CA1 receives inputs from entorhinal cortex through the temporammonic pathway (TA), which has been shown to be important for fear learning and sensitive to stress. Our lab previously showed that NPY reduces TA-evoked synaptic responses, however, the subtype of NPY receptor mediating this effect is not yet known. In this study, we show that Y1 receptors mediate the effects of both exogenous (bath-applied) and endogenously-released NPY in the TA pathway. This is the first demonstration of a Y1 receptor-mediated effect on synaptic function in CA1. Interestingly, chronic cell-type specific overexpression of NPY impairs the sensitivity of the TA pathway to NPY and the Y1 receptor agonist. However, the effect of NPY in the Schaffer collateral (SC) pathway, which is mediate by NPY Y2 receptors, is unaffected by NPY overexpression. There are pathway-specific differences in NPY receptors that modulate NPY’s effects in CA1 and respond differently to NPY overexpression. Our results demonstrating that NPY acts at Y1 receptors in the TA pathway are consistent with the idea that the TA pathway underlies the anxiolytic effects of NPY in CA1.

## Introduction

Neuropeptide Y (NPY), one of the most abundant neuropeptides in the brain (Allen et al., 1983; Reichmann and Holzer, 2016; Smith et al., 2019), has robust neuromodulatory effects in the central nervous system (Colmers et al., 1987, 1985; van den Pol, 2012). NPY mediates a variety of brain functions, such as circuit excitability (Giesbrecht et al., 2010; Silveira et al., 2023), and is implicated in a variety of diseases including epilepsy (Cattaneo et al., 2020). NPY is a stress resilience factor (Reichmann and Holzer, 2016; Sajdyk et al., 2008; Silveira Villarroel et al., 2018) and an important mediator of circuits that regulate mental health (Kupcova et al., 2022; Morales-Medina et al., 2010; Rasmusson and Pineles, 2018; Reichmann and Holzer, 2016). NPY is a major anxiolytic neuropeptide in the brain (Kautz et al., 2017; Nahvi and Sabban, 2020; Wu et al., 2011). In humans, a single nucleotide polymorphism in the promoter region of NPY, rs16147, decreases NPY expression and is correlated with enhanced anxiety traits (Sommer et al., 2010; Zhou et al., 2008). NPY has robust anxiolytic properties (Christiansen et al., 2014; Cohen et al., 2012; Heilig et al., 1989; Karlsson et al., 2005) and reduced NPY is implicated in anxiety disorders, the most common form of mental health disorder (Enman et al., 2015; Kautz et al., 2017; Schmeltzer et al., 2016). NPY is reduced in CSF of patients with post-traumatic stress disorder (PTSD) (Sah et al., 2014) and brain tissue of rodent models of PTSD (Cohen et al., 2012). NPY also has anti-depressive properties (Andriushchenko et al., 2022; Redrobe et al., 2002b, 2002a), and lower NPY levels are found in patients with major depressive disorder (Bale and Doshi, 2023; Guilloux et al., 2012; Hashimoto et al., 1996; Tural and Iosifescu, 2020) and in rodent models of depression (Heilig, 2004; Jiménez-Vasquez et al., 2007, 2000). Multiple antidepressant treatments have been shown to increase NPY levels in rodent models (Bjørnebekk et al., 2010; Husum and Mathé, 2002; Jiménez-Vasquez et al., 2007). NPY itself has been proposed as a potential therapy; in line with this idea, clinical trials have tested intranasal NPY in modulating symptoms for PTSD (Sayed et al., 2018) and depression (Mathé et al., 2020). Despite the fact that NPY’s effects on anxiety and depression have been known for over 30 years (Heilig et al., 1989; Widerlöv et al., 1988), the mechanisms by which NPY modulates synaptic and circuit function to affect mental health are only partially understood.

Anxiety disorders are considered maladaptive forms of learning, and hippocampal dysfunction is implicated in anxiety disorders and PTSD (Çalışkan and Stork, 2019; Cominski et al., 2014; Goosens, 2011). NPY and its receptors are expressed at high levels in hippocampus (Cohen et al., 2012), particularly in the CA1 region of hippocampus (Pickel et al., 1995) that is important for anxiety and fear learning (Engin and Treit, 2007; Engin et al., 2016; Hirsch et al., 2015; Lovett-Barron et al., 2014). Injection of NPY into CA1 of mice is anxiolytic (Smiałowska et al., 2007), and alleviates behavioral symptoms in an animal model of traumatic stress, the Predator Scent Stress model (Cohen et al., 2012). Interestingly, within CA1 there is high expression of NPY dense-core vesicles in *s. lacunosum moleculare* (SLM), the layer of CA1 containing the temporammonic (TA) pathway, a stress-sensitive input pathway (Kallarackal et al., 2013; Kvarta et al., 2015). NPY has long been known to reduce excitatory synaptic transmission of the other main input to CA1, the Schaffer collateral (SC) pathway (Colmers et al., 1988). This effect is through NPY Y2 receptors (Colmers et al., 1991; Greber et al., 1994; McQuiston and Colmers, 1996). Recently, we showed that NPY also inhibits the TA pathway (Li et al., 2017), although the receptor(s) underlying this effect is not yet known.

Injection of NPY into CA1 has anxiolytic effects (Smiałowska et al., 2007) and alleviates anxiety-like behavior in a rodent model of PTSD (Cohen et al., 2012). Both effects are blocked by Y1 receptor antagonism (Cohen et al., 2012; Smiałowska et al., 2007), indicating an essential role for Y1 receptors in CA1 in reducing anxiety. Until now, however, NPY’s only known effects on CA1 synaptic transmission have been via presynaptic Y2 receptors in the SC pathway (Colmers et al., 1991), which mediate NPY’s antiepileptic properties. The location of Y1Rs in CA1 that mediate NPY’s anxiolytic effects is unknown, and effects of Y1Rs on synaptic function in CA1 have yet to be demonstrated. CA1 pyramidal cells express Y1R mRNA (Parker and Herzog, 1998), raising the possibility that Y1Rs in these cells mediate NPY’s effects in the TA pathway. We showed that the NPY dose-response relationship is different between TA and SC synapses (Li et al., 2017), suggesting the receptor type might be different between these two inputs to CA1. Consistent with this, immunostaining for Y2Rs shows no expression in CA1 SLM (Hörmer et al., 2018). Therefore, we hypothesize that Y1 receptors mediates NPY’s effects at TA synapses.

Interestingly, a previous study showed that overexpression of NPY in rats had a reduction in Y1 receptor binding in hippocampus, but not Y2 receptor binding (Thorsell et al., 2000). We previously showed that there is impaired sensitivity to exogenous NPY application at TA-CA1 synapses in slices from mice that overexpress NPY entopically (Corder et al., 2020). Surprisingly, these mice displayed heightened avoidance behavior on the elevated plus maze (Corder et al., 2020), rather than the expected reduction in avoidance behavior seen with NPY injection (Cohen et al., 2012). But, NPY overexpression in these same mice did reduce severity of kainate-induced seizures (Ste Marie et al., 2005), potentially suggesting that there are still functional NPY Y2 receptors. It is not yet known whether NPY overexpression impairs NPY Y1 receptors in the TA pathway, nor whether Y2 receptors in the SC pathway remain functional.

Here we demonstrate that TA-CA1 synapses are modulated by NPY Y1 receptors but not Y2 receptors. In addition, NPY overexpression impairs Y1 receptor function in TA in heterozygous NPY overexpression mice. In contrast, the function of NPY Y2 receptors at SC-CA1 synapses is intact. Together, these results show that there is pathway-specific regulation of the inputs to CA1 by NPY, which are differentially impacted by NPY overexpression.

## Methods

### Animals

All experimental protocols conducted were approved by the Institutional Animal Care and Use Committee at the University of Alabama at Birmingham (APN 20119). All experiments were conducted in accordance to the Guide for the Care and Use of Laboratory Animals adopted by the National Institutes of Health.

Mice were group housed with 3–7 same-sex littermates/cage after they were weaned (p24-p28). Mouse colonies were maintained at 21 ± 2 °C with food and water ad libitum on a 12 hour light/dark cycle. The initial NPY-Tet+/− mouse line was obtained from Jackson Laboratory (B6;129S4-Npytm2Rpa/J; stock no. 007585) (Ste Marie et al., 2005) and maintained on a C57BL/6J background. This mouse line contains a tetracycline regulatory cassette knock-in into the neuropeptide Y gene resulting in overexpression of NPY. Experiments were conducted with age-matched male and female mice from p28-p80 unless otherwise noted. Estrous cycle was not accounted for in females.

### Electrophysiology Recordings

Mice were anesthetized using isoflurane and decapitated with a guillotine. The brain was then rapidly removed and 400 μm thick coronal slices of hippocampus were made with a vibrating microtome (Campden) using standard methods (Li et al., 2017). Slices from ventral hippocampus were collected and CA3 was removed from each slice. Dissection solution was kept ice cold (1–3°C) and contained the following (in mM): 120 NaCl, 3.5 KCl, 0.7 CaCl_2_, 4.0 MgCl_2_, 1.25 NaH_2_PO_4_, 26 NaHCO_3_, and 10 glucose, bubbled with 95% O_2_/5% CO_2_, pH 7.35– 7.45. Slices were allowed to recover at room temperature in dissection solution for >1 hour prior to recording. Recordings were conducted in a submersion recording chamber perfused with external recording solution, a custom artificial cerebrospinal fluid, containing (in mM): 120 NaCl, 3.5 KCl, 2.5 CaCl_2_, 1.3 MgCl_2_, 1.25 NaH_2_PO_4_, 26 NaHCO_3_ and 10 glucose, bubbled with 95% O_2_/5% CO_2_, pH 7.35–7.45. Recordings were conducted at 22–24° C.

Field excitatory postsynaptic potential (fEPSP) were measured in the temporoammonic (TA) pathway in stratum lacunosum-moleculare (SLM) or in the Shaffer collateral (SC) pathway within stratum radiatum (SR). A recording electrode (glass micropipette filled with external recording solution; 2–5 MΩ) and a bipolar tungsten stimulating electrode (FHC, Bowdoinham, ME) or theta glass stimulating electrodes with silver chloride wires were placed in SLM or SR. The synaptic response was measured as the initial slope of the fEPSP. Paired-pulse stimulation was applied using 50 (or 60), 100, 200, 500, and 1000 ms intervals. A 15–20 minute stable baseline was obtained before the onset of each experimental condition by setting the stimulation at the intensity that generated 50–75% of the maximum synaptic response (the largest fEPSP before population spikes are generated). Paired-pulse ratios were calculated as the slope of response 2/slope of response 1.

All electrophysiology experiments were conducted in the presence of the NMDA receptor antagonist, D-2-amino-5-phosphonovalerate (D-APV, 50-100 μM), to prevent long-term potentiation, unless noted otherwise. Neuropeptide Y (1.5 μM) and Leu31-Pro34-NPY (100 nM), the Y1 receptor agonist, were obtained from Bachem or Tocris. PYY, the Y2 receptor agonist (100 nM), BIBP-3226 (Y1 receptor antagonist, 1-2 μM), and BIIE-0246 (Y2 receptor antagonist, 1-2 μM) were obtained from Tocris.

### NPY Release Assay

To test for the effects of endogenously released NPY on synaptic responses, stimulation was compared without (control) and with application of NPY receptor antagonists during stimulation with a physiologically spike train (PST) pattern (Dobrunz and Stevens, 1999; Li et al., 2017; Sun et al., 2018). The PST was based on a pattern of action potential timing previously recorded in vivo from a pyramidal cell in area CA3, provided by Dr. György Buzsáki (New York University, New York, NY) (Isomura et al., 2006). In each PST experimental condition, a pattern of 66 points was presented 3–5 times, separated by a baseline section of 25 points at a slow constant frequency (0.083 Hz). fEPSPs slopes for each recording are normalized by the average slope value of the last 10-15 points of the baseline period and plotted against stimulus number to show short-term plasticity. Between experimental conditions, there was at least a 10-minute washin period for each NPY receptor antagonist before another round of the PST was applied. Stimulus intensity and duration were constant during each experiment.

### ELISA

Mice were anesthetized with isoflurane and euthanized by decapitation and trunk blood was collected and placed into tubes containing heparin, aprotinin, and DPP-IV inhibitor for NPY levels or in tubes containing only heparin for corticosterone levels. Blood samples were spun at 3,000 RPM for 15 min at 4°C in a microcentrifuge. Plasma was collected and frozen with dry ice. Samples were stored at −20°C. Neuropeptide Y levels were measured using commercially available ELISA kits (EMD Millipore, EZHNPY-25K).

Neuropeptide Y levels in tissue were measured with commercially available ELISA kit (EMD Millipore, EZHNPY-25 K). Mice were anesthetized with isoflurane and euthanized by decapitation. Hippocampus was collected and flash frozen with dry ice. Samples were homogenized in NP-40 homogenization buffer (10 mM Tris pH 7.5,10 mM NaCl, 3 mM MgCl_2_·6H_2_O, 1 mM EDTA, 0.5M of 10% NP-40) and centrifuged at 16,000 ×g for 10 min at 4 °C. Supernatant was collected and stored at −80 °C. A Bradford Assay was conducted to ensure uniform total protein level loading (1 μg/ well) into the ELISA assay.

### Statistics

Data are reported as mean ± S.E.M. Data were tested for normal distribution and variance to determine whether to use parametric or nonparametric statistical tests. Data sets in which the means between multiple groups, such as comparing hippocampal NPY levels across genotype, a One-Way ANOVA was used with a LSD post hoc test. For pharmacology experiments in which a baseline period was compared to a single drug treatment, a paired T-test was used. If multiple drugs were used or if a dose response experiment was performed, a repeated measures ANOVA was used with a LSD post-hoc test to determine significance across groups. test was used to determine statistical outliers. If the data did not exhibit equal variance, a Welch correction was used.

## Results

### Effects of pharmacological application of NPY are reversed in SC, but not TA evoked fEPSPs

We previously showed that the effects of NPY bath application differ between SC-evoked versus TA-evoked synaptic responses recorded in CA1. As the effects of NPY on TA stimulated fEPSPs are smaller than the effects on SC-stimulated fEPSPs, we reconfirmed that NPY reduces the synaptic response in the TA pathway (Figure 1) by comparing vehicle to 1 μM NPY treated slices. As observed in Figure 1D, only the application of NPY reduced TA fEPSP responses. Therefore, the effect is not due to synaptic rundown of TA-fEPSPs. Additionally, a concern had arisen that NPY’s effect in the TA pathway might have been partially due to contamination from the SC pathway. Therefore, in a subset of experiments, the TA pathway was confirmed with a 1-2 μM application of DCG-IV. This low dose of DCG-IV basically abolishes TA fEPSPs with no effects on SC fEPSPs (data not shown). In these chemically identified TA fEPSPs, NPY still reduced TA-induced synaptic response (data not shown).

**Figure 1:**
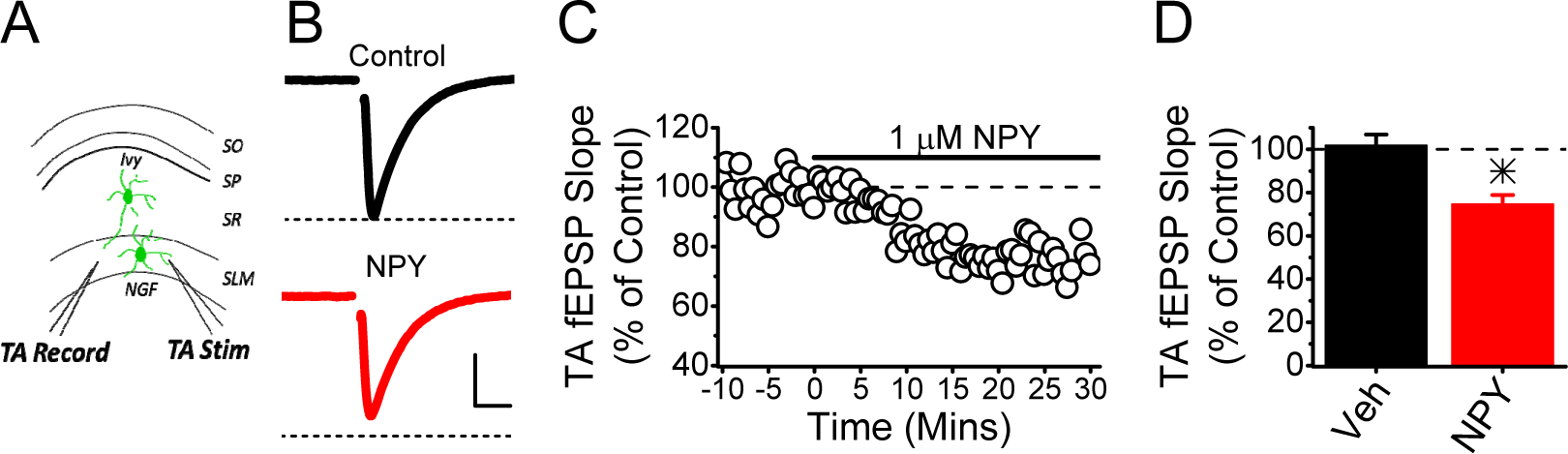
Bath-applied NPY reduces TA fEPSPs. **A)** Schematic showing field potential recordings from SLM in response to Temporoammonic stimulation (TA) **B)** Example traces of TA-evoked fEPSPs with (red) and without (black) the application of 1 µM NPY. Scale Bars: 0.4 mV, 10 ms **C)** Example experiment showing the time course of the effect of NPY on TA-evoked fEPSPs. **D)** Exogenous NPY decreases the synaptic response size in the TA pathway, whereas vehicle has no effect (t-test, p <0.05, n = vehicle 4 slices / 4 animals, NPY 8 slices / 8 animals). (*p <0.05 vehicle vs treatment).

It is unknown the specific receptor underlying NPY’s effect on the TA pathway. As NPY Y1 and Y2 receptors expression has been observed in hippocampus (Hörmer et al., 2018; Méndez-Couz et al., 2021; Parker and Herzog, 1999), we first tested whether either Y1 or Y2 receptor antagonists could reverse the effects of bath applied NPY in the TA pathway. Bath application of NPY alone caused a reduction in the response of TA fEPSPs to approximately 60% of control (Figure 2B, C). The effect was not reversed by either the Y1 receptor antagonist (BIBP3226, 2 µM), or subsequent addition of the Y2 receptor antagonist (BIIE0246, 2 µM, Figure 2B, C). In SC-evoked responses, we found that NPY suppresses the fEPSP to approximately 40% of control, an effect that was not altered by application of the Y1 receptor antagonist BIBP3226 alone (Figure 2E, F). However, subsequent addition of the Y2 receptor antagonist BIIE0246 reversed the effects of NPY, consistent with previous studies showing that Y2 receptors mediate NPY’s effects in the SC pathway (Figure 2 E, F) (Colmers et al., 1991; Greber et al., 1994; McQuiston and Colmers, 1996). Together, the data indicate that SC and TA differ in their responsiveness to exogenously applied NPY, but do not indicate which receptor mediates NPY’s effects in TA.

**Figure 2:**
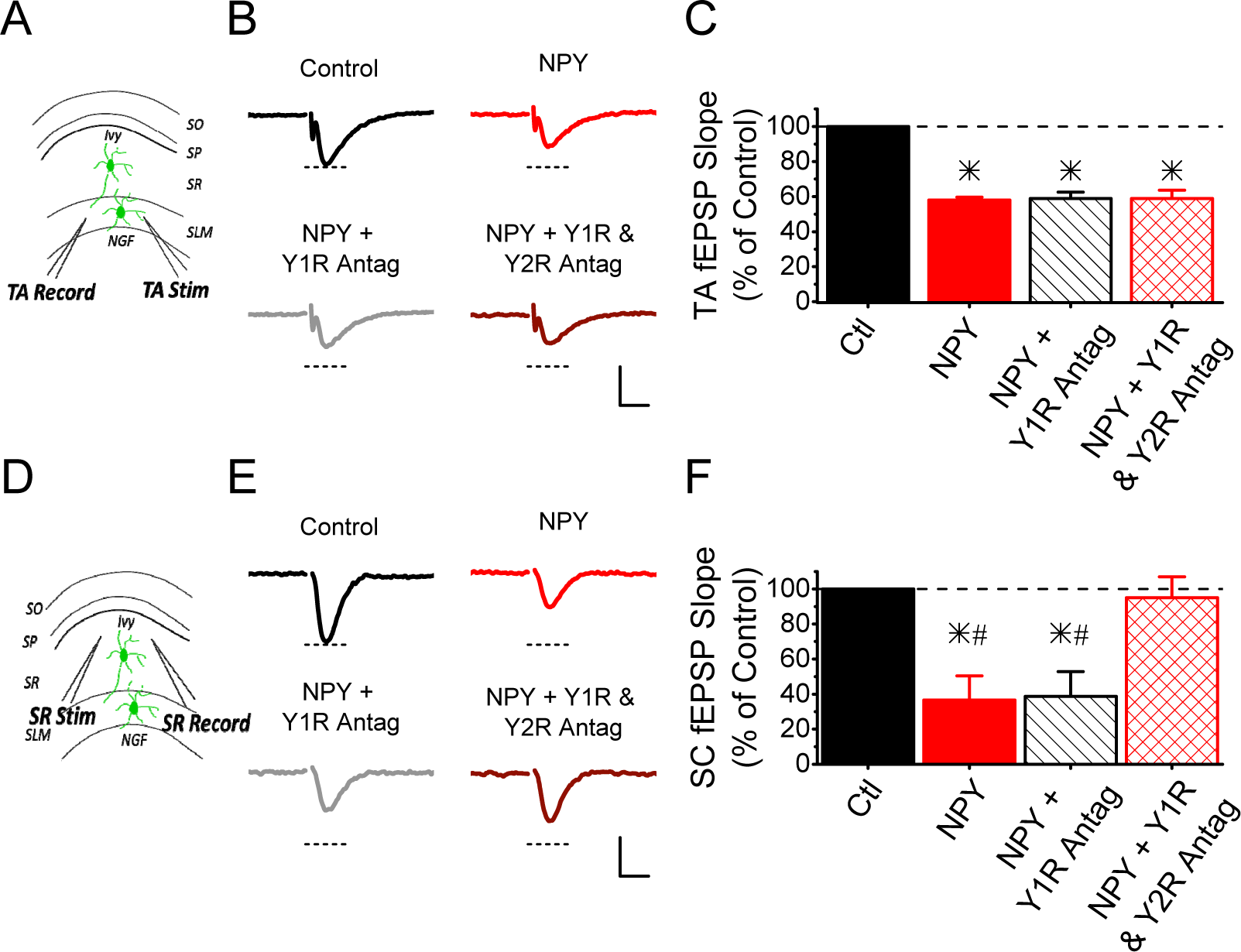
NPY’s effect in the TA pathway is not reversed by Y1 or Y2 antagonists. **A)** Schematic showing field potential recordings from SLM in response to Temporoammonic stimulation (TA) **B)** Example traces of TA-evoked fEPSPs with (red) and without (black) the application of 1.5 µM NPY, then with the addition of the Y1 receptor antagonist (gray, 2 µM BIBP3226), and finally with the application of the Y2 receptor antagonist (maroon, 2 µM BIIE0246). Scale Bars: 0.2 mV, 10 ms. **C)** Exogenous NPY decreases synaptic response size in the TA pathway. The effect of NPY on the synaptic response is not reversible with either NPY Y1 receptor (BIBP3226) or Y2 receptor (BIIE0246) antagonist (Repeated Measures ANOVA main drug effect, F_(3,6)_ =74.51, p<0.05; n = 3 slices / 3 animals). **D)** Schematic showing field potential recordings from s. radiatum in response to Schaffer collateral (SC) stimulation. **E)** Example traces of SC-evoked fEPSPs with (red) and without (black) the application of 1.5 µM NPY, then with the addition of the Y1 receptor antagonist (gray, 2 µM BIBP3226), and finally with the application of the Y2 receptor antagonist (maroon, 2 µM BIIE0246). Scale Bars: 0.2 mV, 10 ms. **F)** Exogenous NPY decreases synaptic response size in the SC pathway. The effect of NPY can be reversed by the application of the NPY Y2 receptor antagonist, but not by the NPY Y1 receptor antagonist (Repeated Measures ANOVA main effect of drug, F_(3,9)_ = 12.45, p < 0.05; n = 4 slices / 4 animals). (*p <0.05 treatment vs control) (#p <0.05 treatment vs both antagonists).

### Y2R agonism mimics NPY’s effects at SC synapses but not TA synapses

To better understand the receptors that are mediating NPY’s effects at TA synapses, we used Y2 receptor specific agonist, PYY_3-36_, to determine if activation of Y2 receptors can mimic the effects of NPY. We find that 100 nM PYY_3-36_ does not affect TA-CA1 synaptically evoked fEPSPs (Figure 3B), suggesting that the effects of NPY on TA synapses are not mediated by Y2 receptors. Additionally, we show that application of PYY_3-36_ does not alter the paired pulse ratio of TA-evoked fEPSPs at the 50 millisecond interval (Figure 3C). As a control, we bath applied 100 nM PYY_3-36_ and recorded SC-evoked fEPSPs in CA1 and found that it mimicked NPY’s effects, as seen by a reduction to 40% of the control response size (Figure 3E), as previously reported (Colmers et al., 1987; El Bahh et al., 2005, 2002). Y2 receptor antagonism by 1µM BIIE02246 partially reversed PYY_3-36_’s effects at SC-CA1 synapses (Figure 3E). Additionally, we find that PYY_3-36_ application to SC-evoked fEPSPs enhanced the paired pulse ratio at the 50 millisecond interval (Figure 3F), agreeing with previous studies showing that NPY Y2 receptors in the SC pathway are presynaptic (Colmers et al., 1991). Together, the data indicate that Y2 receptors do not affect TA-CA1 synaptic transmission.

**Figure 3:**
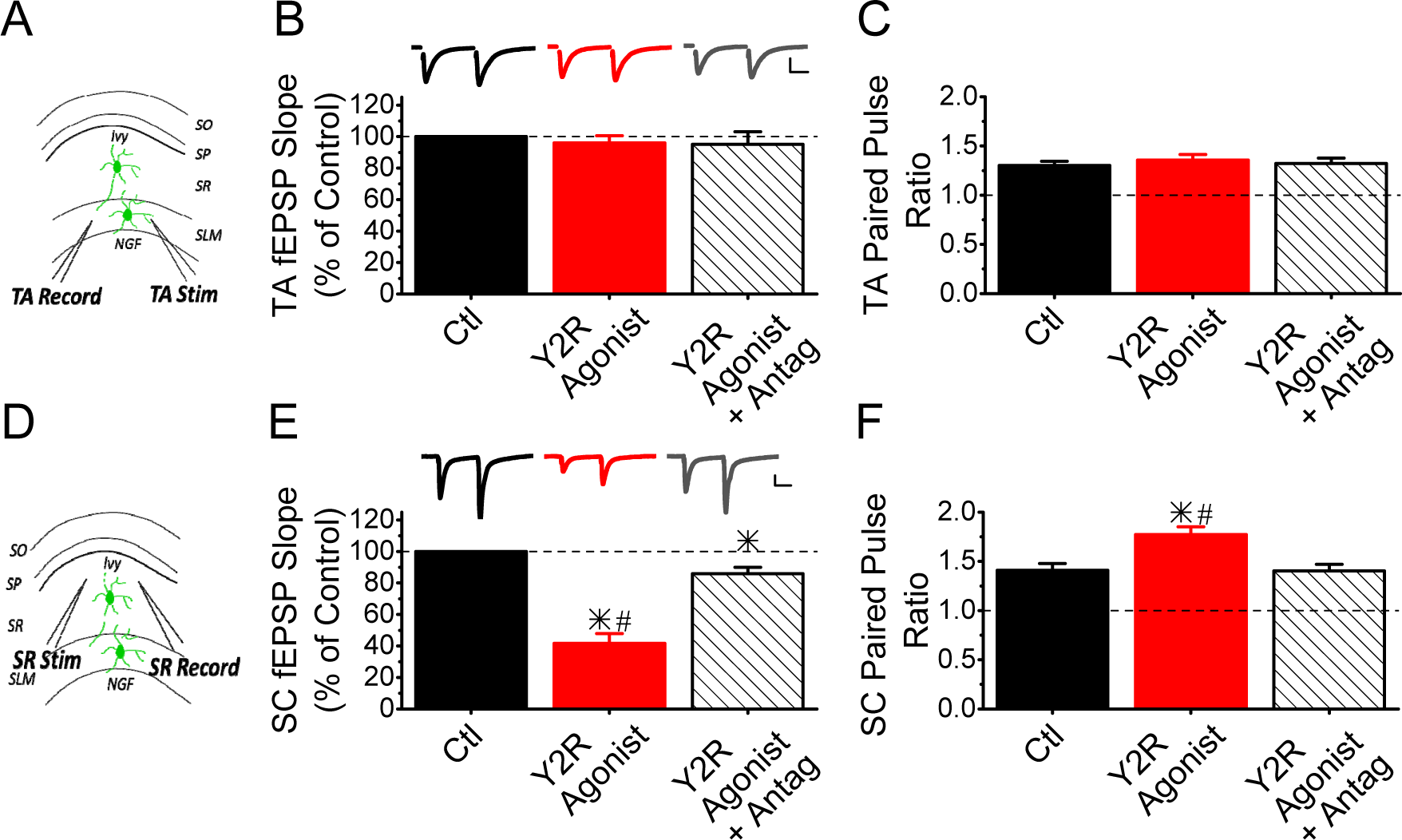
Activation of NPY Y2 receptors does not reduce TA fEPSPs. **A)** Schematic showing field potential recordings from SLM in response to Temporoammonic stimulation (TA) **B)** Exogenous PYY 3-36 (100 nM), the NPY Y2 receptor agonist, had no effect on the fEPSPs evoked by stimulating the TA pathway. Additionally, the NPY Y2 receptor antagonist did not affect the TA evoked fEPSPs. (Repeated Measures ANOVA, F_(2,8)_ = 0.41, p = 0.68; n = 5 slices / 5 animals). Inset: Example traces of TA evoked fEPSPs during a control period (black), in the presence of PYY 3-36 (red), and with the addition of 1 µM BIIE0246 (dark gray). Scale Bars: 0.2 mV, 20 ms. **C)** The paired-pulse ratio at a 50 ms interval was unchanged by the presence of PYY 3-36 and BIIE0246 for TA synapses. (Repeated Measures ANOVA, F_(2,8)_ = 0.62, p = 0.56; n = 5 slices / 5 animals). **D)** Schematic showing field potential recordings from s. radiatum in response to Schaffer collateral (SC) stimulation. **E)** Activation of NPY Y2 receptors with PYY 3-36 reduced fEPSPs evoked by stimulating the SC pathway. The effect was partially reversed by the application of the NPY Y2 receptor antagonist (BIIE0246). (Repeated Measures ANOVA main effect of drug, F_(2,12)_ = 64.66, p < 0.05; n = 7 slices / 7 animals). Inset: Example traces of SC evoked fEPSPs during a control period (black), in the presence of PYY 3-36 (red), and with the addition of BIIE0246 (dark gray). Scale Bars: 0.2 mV, 20 ms. **F)** Exogenous PYY 3-36 enhances the paired-pulse ratio of SC fEPSPs at the 50 ms interval, and the effect is reversed by the application of BIIE0246. (Repeated Measures ANOVA main effect of drug, F_(2,12)_ = 41.70, p < 0.05; n = 7 slices / 7 animals). (*p <0.05 treatment vs control; #p<0.05 agonist vs antagonist).

### Y1R agonism reduces evoked fEPSPs from TA-CA1 synapses

To determine if Y1 receptors could be mediating NPY’s effects at TA synapses, we tested the effects of bath application of the Y1 receptor preferring agonist Leu^31-^Pro^34-^NPY (L-P-NPY) (McQuiston et al., 1996). We performed a dose response experiment and found that 100 nM Leu^31^Pro^34^NPY resulted in a maximum reduction of the TA synaptic response size (data not shown). Bath application of 100 nM Leu^31-^Pro^34-^NPY to TA-evoked fEPSPs in CA1 resulted in reduction to about 70% of the control response size (Figure 4A), similar to what was observed with application of NPY. This is the first indication that Y1 receptor activation suppresses the excitatory synaptic response in the TA pathway. We also find that 100 nM Leu^31^-Pro^34^-NPY at TA-CA1 synapses does not affect paired pulse ratio at the 50 millisecond interval (Figure 4B).

**Figure 4.**
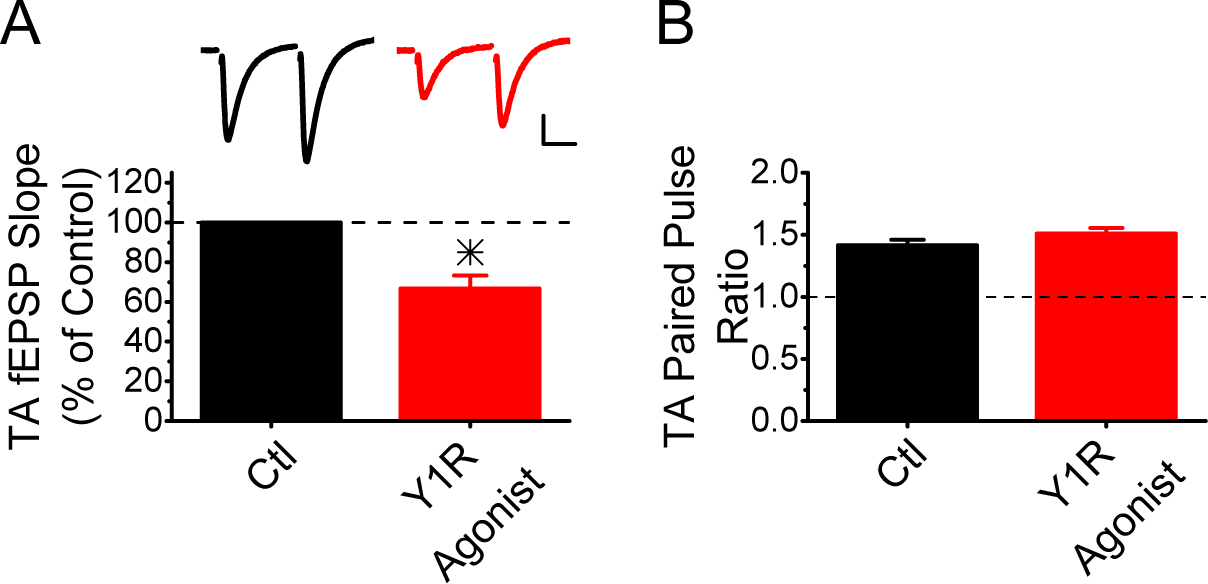
Activation of Y1 receptors reduces TA fEPSPs. **A)** Exogenous [Leu31,Pro34]-NPY (100nM), the NPY Y1 receptor agonist, reduced TA-evoked fEPSPs. (paired t-test, p <0.05; n = 6 slices / 6 animals). **B)** Exogenous [Leu31,Pro34]-NPY had no effect on the paired pulse ratio of TA-evoked fEPSPs (paired t-test, p =0.18; n = 6 slices / 6 animals). (*p <0.05 treatment vs control).

### Effects of endogenous NPY release are blocked by Y1 but not Y2 receptor antagonism in TA

We next wanted to determine if Y1 receptors also mediate the effects of endogenously released NPY in the TA pathway. Our lab has previously shown that NPY is released by electrical stimulation of TA synapses with a physiological spike train (PST), which suppresses short-term plasticity (Li et al., 2017). This effect was unmasked by application of a combination of Y1 and Y2 receptor antagonists (BIBP3226 + BIIE0246) (Li et al., 2017). Here, we use this same stimulation protocol to induce endogenous NPY release at TA-CA1 synapses, but sequentially applied BIBP3226 and then BIIE0246, or vice versa, to determine which receptor may be mediating the effects of endogenously released NPY in the TA pathway.

Stimulation with the PST (3-5 repetitions of a 66 point PST with a 25 point control period for each repetition) causes a mixture of short-term facilitation and short-term depression, as shown in Figure 5B. Blocking Y1 receptors with bath application of the Y1 receptor antagonist BIBP3226 alone was sufficient to increase the amount of facilitation during subsequent stimulation with the same PST pattern (Figure 5B, red open circles), with almost no further enhancement with the subsequent application of BIIE0246 (Figure 5C, gray closed circles). This is quantified in Figure 5D, which showed that blocking Y1 receptors increased the average response to the PST. Surprisingly, group data show that blocking Y2 receptors caused an additional small yet statistically significant increase in the amount of short-term facilitation. Conversely, we find that application of BIIE0246 alone had no effect on short-term plasticity in response to the PST, but subsequent application of BIBP3226 did result in enhanced short-term facilitation of TA-CA1 fEPSPs (Figure 5E). Both of these experiments indicate that Y1 receptors modulate the effects of endogenously released NPY in TA. There do not appear to be Y2 receptors acting as autoreceptors to limit NPY’s release, because blocking Y2 receptors alone had no effect on short-term plasticity in response to the PST. Together, the data indicate that endogenously released NPY signals through Y1 receptors at TA-CA1 synapses.

**Figure 5.**
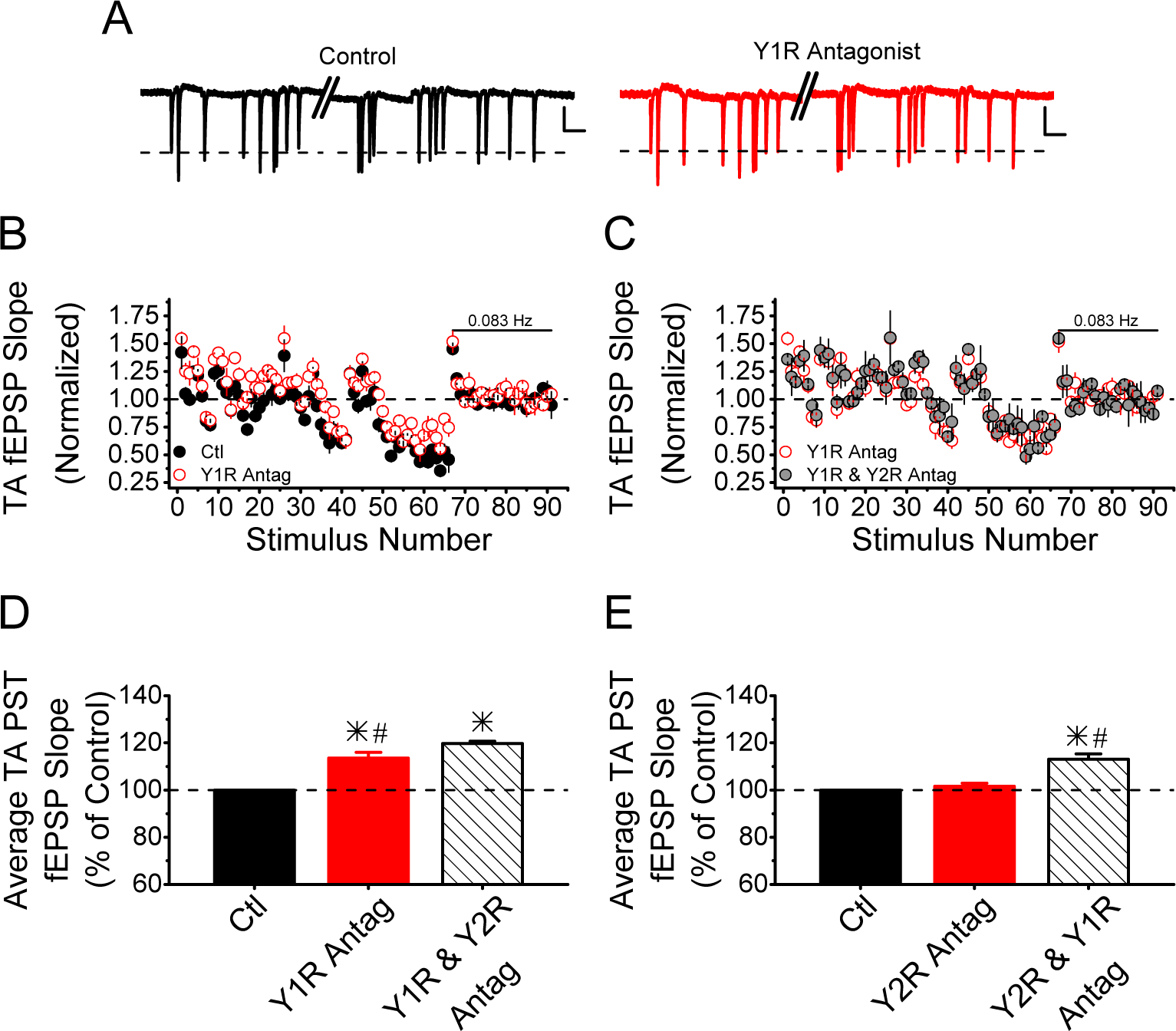
Endogenously released NPY in SLM is mediated by the NPY Y1 receptor. **A)** Example fEPSP traces during a portion of the physiological spike train (PST) pattern in response to TA stimulation with (red) and without (black) application of the Y1 receptor antagonist, 1 μM BIBP3226. The fEPSP traces show the TA synaptic response to the first 21 points in the PST pattern. To improve visibility of the synaptic response to the PST 3.5 seconds was excluded as no stimulation occurred during this time. The break in the trace is indicated by two diagonal lines. The dashed line indicates the average fEPSP peak amplitude during the baseline period of the PST. Scale Bar: 0.2 mV, 250 ms. **B-E)** The PST pattern is a variable pattern of stimulation and has interstimulus intervals ranging from 30 ms to 23.6 s. There are a total of 66 points in the PST pattern. Each PST pattern is repeated three to five times during the control and each successive application of a NPY receptor antagonists. Responses are normalized to a 25-point baseline period (0.083 Hz constant frequency) applied at the end of each PST repetition. **B)** An example experiment showing the mixture of short-term plasticity observed at TA synapses onto CA1 pyramidal cells during the PST pattern with (red open circle) and without (black circle) the NPY Y1 receptor antagonist. Blocking NPY Y1 receptor shifts the short-term plasticity values upwards during the PST pattern, but not during the baseline period, suggesting endogenous release of NPY. Data is plotted as normalized fEPSP slope vs. stimulus number during the PST pattern (first 66 points), normalized by the average slope size during the baseline period at the end (0.083 Hz constant frequency, last 25 points). **C)** An example experiment showing that the addition of the NPY Y2 receptor antagonist (BIIE0246) has little effect on enhancing the short-term plasticity further when compared to BIBP3226 alone. This data is a continuation of the example experiment in C. **D)** Blocking NPY Y1 receptors increases the amount of short-term facilitation at TA synapses onto CA1 pyramidal cells during PST stimulation, causing an ∼ 12% increase in the average normalized fEPSP slope. Adding the NPY Y2 receptor antagonist did slightly enhance the effect further. (Repeated Measures ANOVA main effect of drug, F_(2,6)_ = 66.03, p < 0.05; n = 4 slices / 4 animals). **E)** Blocking NPY Y2 receptors first had no effect on the short-term plasticity at TA synapses onto CA1 pyramidal cells during PST stimulation. Adding the NPY Y1 receptor antagonist enhanced the amount of short-term facilitation, causing an ∼ 13% increase in the average normalized fEPSP slope (Repeated Measures ANOVA main effect of drug, F_(2,4)_ = 33.22, p < 0.05; n = 3 slices / 3 animals). (*p <0.05 treatment vs control; #p<0.05 single antagonist vs both antagonist).

### Mice transgenically overexpressing NPY have Y1R loss function at TA-CA1 synapses but functional Y2R at SC-CA1 synapses

NPY^tet^ mice overexpress NPY in a cell-type specific manner (Ste Marie et al., 2005). Our lab previously identified that homozygous mice (NPY^tet/tet^) have impaired bath-applied NPY sensitivity at TA-CA1 synapses (Corder et al., 2020) and a slight increase in anxiety. Based on our previous data and the fact the loss of NPY Y1 receptors enhances anxiety (Bertocchi et al., 2011; Karl et al., 2006), overexpression of NPY causes impairment of NPY Y1 receptors. However, the NPY^tet/tet^ was able to reduce the seizure severity (Ste Marie et al., 2005), and since NPY Y2 receptors mediates the ‘anti-seizure’ effect (El Bahh et al., 2002; Ledri et al., 2015), it is possible that NPY Y2 receptors are still functional in these mice. Additionally, it is unknown how different levels of NPY overexpression can affect NPY receptors.

Therefore, we wanted to determine if Y2 receptors at SC-CA1 synapses are impervious to effects of chronic overexpression of NPY, and what level of NPY expression causes impairment of Y1 receptor function at TA synapses. First, we wanted to determine the level of NPY expression in NPY^tet/wt^. We show that heterozygous mice (NPY^tet/wt^) have an intermediary level of NPY expression in hippocampus relative to wildtype (NPY^wt/wt^) and homozygous (NPY^tet/tet^) mice (Figure 6A).

**Figure 6:**
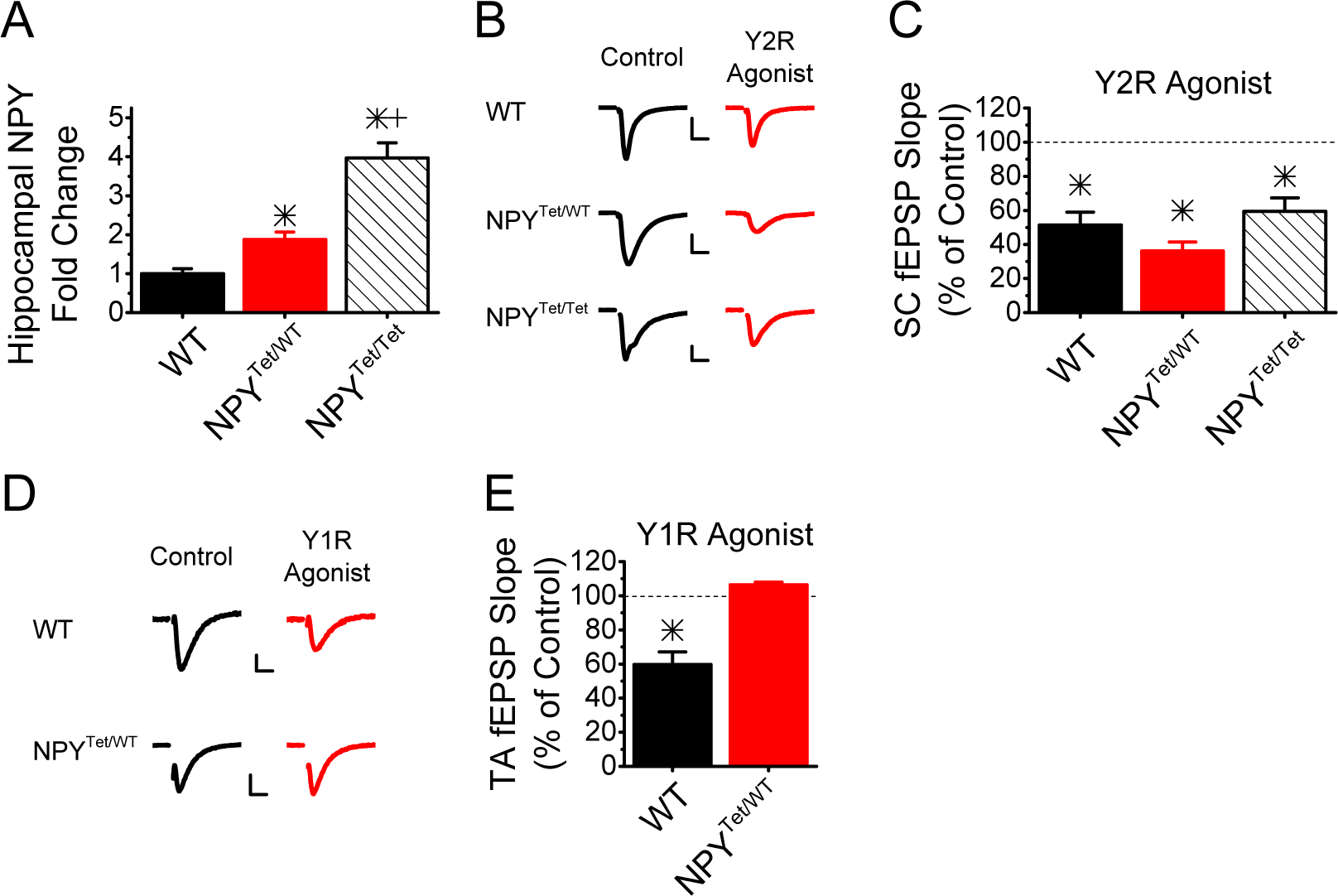
Chronic overexpression of NPY has no effect on SC Y2 receptors but causes impairment of TA Y1 receptors. **A)** NPY^tet/wt^ have a 1.5-fold overexpression of NPY in hippocampus, while NPY^tet/tet^ mice have a 3-fold overexpression relative to wild type mice (One way ANOVA F_(2,18)_=34.51, p<0.05; n=6 animals per genotype) (*p<0.05 genotype vs WT; +p<0.05 Het vs Hom) **B)** *Left inset* Example SC-evoked fEPSP control traces (black) from wild type (top), NPY^tet/wt^ (middle), and NPY^tet/tet^ (bottom) mice. *Right inset* Example SC-evoked fEPSP traces in the presence of 100nM PYY3-36 (red) from each genotype. **C)** SC-evoked fEPSPs are reduced by PYY in slices from all genotypes to a similar extent. (Two-way ANOVA main effect of drug treatment, F_(1,48)_=197.05, p<0.05; n=10, 10, 7 animals) (*p <0.05 treatment vs control). No effect of genotype was observed on activation of NPY Y2 receptors (Two-way ANOVA main effect of genotype, F_(2,48)_=1.25, p=0.30; n=10, 10, 7 animals). **D)** *Left inset* Example TA-evoked fEPSP control traces from wild type (top) and NPY^tet/wt^ (bottom) mice. *Right inset* Example TA-evoked fEPSP traces in the presence of 100 nM [Leu31,Pro34]-NPY from wild type (top) and NPY^tet/wt^ (bottom) mice. **E)** TA-evoked fEPSP responses are reduced in slices from wild type mice (n=5 slices/5 animals) with the application of [Leu31,Pro34]-NPY, but not in slices from NPY^tet/wt^ (n=3 slices/3 animals), revealing a drug-genotype interaction (Two-way ANOVA main effect of genotype x drug treatment, F_(1,12)_=18.41, p<0.05) (*p <0.05 treatment vs control; # WT vs Het).

To test if Y2 receptors at SC are still functional with NPY overexpression, we applied the Y2 receptor agonist, PYY_3-36_ (100 nM), to slices from NPY^wt/wt^, NPY^wt/tet^, and NPY^tet/tet^ mice. We found the typical reduction to about 40% of control synaptic responses was intact in all three genotypes (Figure 6 B and C). Therefore, showing that the NPY Y2 receptors are still functional even when NPY is overexpressed by almost 3-fold.

To determine if Y1 receptor function was intact at TA-CA1 synapses in heterozygous mice (NPY^tet/wt^), we bath applied the Y1 receptor agonist Leu^31-^Pro^34-^NPY (100 nM) to slices of NPY^wt/wt^ and NPY^tet/wt^ mice. We found that the typical reduction to 60% of control responses in wildtype was not evident in slices from NPY^tet/wt^ mice (Figure 6 D and E). This shows that that NPY receptors are impaired in the TA pathway in heterozygous NPY overexpression mice, as has previously been shown for homozygous mice (Corder et al., 2020). Together, our data indicate that transgenic overexpression of NPY differentially affects synaptic inputs to area CA1 by impairing Y1 receptor function at TA, while Y2 receptor sensitivity remains intact.

## Discussion

Here we demonstrate for the first time that the effects of NPY at TA-CA1 synapses are mediated by Y1 receptors. This was observed with both bath-applied Y1 receptor agonist and endogenously released NPY. This is in contrast to SC-CA1 synapses, which are known to be mediated by Y2 receptors and not Y1 receptors (Colmers et al., 1991; Greber et al., 1994; McQuiston and Colmers, 1996). Additionally, we observe that chronic entopic overexpression of NPY has differing consequences on hippocampal area CA1 depending on the synaptic input and/or receptor type. Specifically, we find that TA-CA1 synapses have impaired Y1 receptor function at an intermediary level of NPY overexpression observed in NPY^tet/wt^ mice, whereas even the higher levels of NPY overexpression in NPY^tet/tet^ mice did not alter Y2 receptor function at SC-CA1 synapses.

These experiments reveal for the first time an effect of Y1 receptor activation on synaptic transmission in CA1. Previous studies showed that injection of NPY into CA1 increases time spent in the open arms of the elevated plus maze (Cohen et al., 2012; Smiałowska et al., 2007). The effect was blocked by injection of a Y1 receptor antagonist (Cohen et al., 2012; Smiałowska et al., 2007), indicating an essential role for Y1 receptors in CA1 for the anxiolytic response. The perforant pathway input to CA1, which we and others refer to as the temporoammonic (TA) pathway, is particularly sensitive to stress (Cai et al., 2013; Kallarackal et al., 2013; Li et al., 2017) and carries aversive sensory information (Lovett-Barron et al., 2014). Therefore, this pathway could be an important mediator for the anxiolytic response produced by hippocampal CA1. Consistent with this, our lab demonstrated that traumatic stress impairs the release of endogenous NPY in the TA pathway. In addition, traumatic stress caused an increase in avoidance behavior on the elevated plus maze (Li et al., 2017). Therefore, it is tempting to speculate that loss of Y1 receptor-mediated signaling in the TA pathway of CA1 could contribute to the increased anxiety-like behavior in mice after traumatic stress. Furthermore, the impairment of Y1 receptors in TA in the NPY overexpression mice may be a contributing factor to enhanced avoidance behavior in these mice (Corder et al., 2020). Together, these results are consistent with the idea that Y1 receptor function in the TA pathway underlies the anxiolytic effects of NPY in CA1 in rodents.

Our results show pathway-specific differences in the NPY receptors that modulate NPY’s effects on CA1 pyramidal cell inputs. Y1 receptors have a higher affinity for NPY than Y2 receptors (Pronchuk et al., 2002). As a result, they are likely to respond better to physiological (non-pathological) levels of NPY release. Consistent with this, we previously demonstrated that physiologically relevant stimulation of the TA pathway alone resulted in detectable NPY release, whereas stimulation of the SC pathway alone did not (Li et al., 2017). Instead, a combination of SC and TA stimulation was needed to trigger NPY release that modulated the SC pathway (Li et al., 2017). The lower affinity of the Y2 receptors in the SC pathway is consistent with their role in suppressing epileptic activity (Dum et al., 2017; El Bahh et al., 2002; Furtinger et al., 2001; Klapstein and Colmers, 1997; Ledri et al., 2015). The high-frequency burst firing that occurs during epileptic activity is likely to cause a high level of NPY release, reaching concentrations necessary to activate Y2 receptors. While the function of Y1 receptors in the TA pathway is only beginning to be uncovered, the pathway-specific expression of high (Y1) and low (Y2) affinity NPY receptor subtypes enables NPY to differentially modulate the two inputs to CA1.

Importantly, high-affinity NPY Y1 receptors are susceptible to rapid desensitization and internalization following agonist exposure (Gicquiaux et al., 2002; Holliday et al., 2005; Kempf et al., 2015). Consistent with this, we previously showed that in the TA pathway of homozygous NPY ^tet/tet^ mice (Corder et al., 2020), chronic NPY overexpression impaired the function of NPY receptors, which we now know are Y1 receptors. Furthermore, there is also impairment of TA Y1 receptor function in heterozygous NPY overexpression mice, indicating that even more moderate levels of NPY enhancement can be detrimental when chronically elevated. This could be caused by receptor internalization, as has been shown for Y1 receptors in rodent nerve injury models (Marvizon et al., 2019; Nelson and Taylor, 2021). Loss of Y1 receptor function could be caused by downregulation, as prolonged ligand exposure has often been shown to lead to GPCR downregulation (Black et al., 2016). There is reduced Y1 receptor binding in hippocampus of rats overexpressing NPY (Thorsell et al., 2000), consistent with downregulation or internalization of Y1Rs. There could also be uncoupling of Y1Rs from their G proteins, which would lead to impaired GPCR function (Ma et al., 2021). In contrast, Y2 receptor function in the SC-CA1 pathway is intact in both the NPY^tet/wt^ and NPY^tet/tet^ mice, similar to that of wildtype controls. This finding indicates that the Y2 receptors may be less likely to desensitize or downregulate even in the presence of high NPY levels, enabling them to function during pathological conditions.

A previous study showed that NPY overexpression protects against kainate-induced seizures in the NPY ^tet/tet^ mice (Ste Marie et al., 2005), supporting the idea of intact Y2 receptor function. However, they found no effect on feeding behavior in these mice (Ste Marie et al., 2005), which is mediated in part by Y1 receptors (Kalra et al., 1991), indicating that Y1 receptors may also be impaired in other brain regions, such as the hypothalamus. It is likely that other brain regions with high levels of Y1 receptors, such as amygdala (Parker and Herzog, 1999), could also have impaired Y1 receptors. If so, loss of Y1R function in brain regions other than CA1 might also contribute to the mild increase in anxiety-like behavior previously observed in the NPY overexpression mice (Corder et al., 2020). Together, we find that mice that overexpress NPY have pathway- and receptor-specific loss of function, and that Y1 receptors (anti-anxiety) may be more susceptible to downregulation by increased NPY than Y2 receptors (anti-epileptic). Impairments in Y1R function by long-term increases in NPY may limit NPY’s potential benefit as a chronic therapeutic agent.

Unlike Y2 receptors, Y1 receptors are typically postsynaptic (Kopp et al., 2002; Molosh et al., 2013), but presynaptic Y1 receptors are also found (Kopp et al., 2002). Here we saw no significant effect on the paired-pulse ratio of TA synapses in response to the Y1 receptor agonist despite its reducing the first pulse fEPSP response size. This suggests that the Y1 receptors are postsynaptic at TA synapses in CA1. In contrast, our previous study saw increased paired-pulse ratio at TA synapses with bath application of NPY (Li et al., 2017), suggesting a possible presynaptic effect. However, the paired-pulse ratio is an indirect indicator of the location of synaptic changes; although changes in paired-pulse ratio are often caused by presynaptic changes in initial release probability (Zucker et al., 1991), this is not always the case (Sippy et al., 2003; Walters et al., 2014). Synaptosomes prepared from whole hippocampus, dentate gyrus, and CA3 show inhibition of glutamate release via activation of Y1 as well as Y2 receptors, suggesting that Y1 receptors can be located presynaptically in hippocampus (Silva et al., 2001). In contrast, Y1 receptor activation did not inhibit glutamate release in CA1 derived synaptosomes, which were inhibited by Y2 receptor activation (Silva et al., 2001), suggesting that there are not presynaptic Y1 receptors in CA1. It is therefore not yet clear whether the Y1 receptors are acting pre – or postsynaptically at the TA pathway.

Y1 receptors, like Y2 receptors, act through G_i/o_ coupled signaling cascades, resulting in reduced PKA activity (Molosh et al., 2013; Pleil et al., 2015). Y1 receptors have been shown to act through a variety of mechanisms, including postsynaptic effects through I_h_, GIRK, or dendritic calcium channels, and presynaptic effects through inhibition of calcium channels (Giesbrecht et al., 2010; Hamilton et al., 2013; McQuiston et al., 1996; Molosh et al., 2013). Y1 receptors are abundantly expressed on granule cells in dentate gyrus, (Sperk et al., 2007), but they do not modulate excitatory synaptic transmission in hippocampal slices (McQuiston et al., 1996). Instead, Y1 receptors inhibit depolarization induced calcium influx, measured in the granule cell soma, through effects on N-type calcium channels (McQuiston et al., 1996). In amygdala, Y1 receptors causes hyperpolarization of pyramidal neurons through suppression of Ih currents (Giesbrecht et al., 2010). Interestingly the distal dendrites of CA1 where the TA pathway synapses are formed are highly enriched in HCN1 channels that underlie the Ih current (Lörincz et al., 2002). More work is needed to determine if the effects of Y1 receptors at TA synapses are through modulation of dendritic Ih.

While the Y1 receptor antagonist (BIBP3226) used in this study has minimal affinity to other NPY receptors (Doods et al., 1996), the Y1-preferring agonist used, Leu^31^-Pro^34^-NPY, also binds to Y5 receptors with low affinity ((Pronchuk et al., 2002). Because the K_i_ is 10-fold higher for Y5 relative to Y1 receptors ((Pronchuk et al., 2002), the concentration of agonist used for our experiments should activate Y1Rs and not Y5Rs. It is important to note that intracerebroventricular (ICV) infusion of Y5 receptor antagonist does not influence anxiety behavior and neither does Y5 receptor overexpression, although a mild hyperactive phenotype had been observed (Kask et al., 2001; Olesen et al., 2012). To date, Y5 receptor signaling remains unclear in the context of hippocampal circuit function, and our experiments do not rule out a possible role for Y5Rs at TA synapses. Some brain regions contain neurons that coexpress both Y1 and Y5 receptors (Fekete et al., 2002; Gehlert et al., 2007; Longo et al., 2015, 2014; Wolak et al., 2003), and Y1 and Y5 receptors can form heterodimers with altered agonist and antagonist properties (Gehlert et al., 2007). More work is necessary to understand how Y1 and Y5 signaling mechanisms converge and if this is a phenomenon observed within the hippocampus. Lastly, both male and female mice were used in this study, although estrus cycle was not monitored. Because estrogen can regulate NPY (Ledoux et al., 2009) and Y1 receptor expression (Hill et al., 2004) it would be of interest to determine if phase of estrus cycle in female mice also affects NPY receptor expression and downstream NPY receptor signaling in the TA pathway.

Overall, we provide the first demonstration of TA-CA1 synapses being sensitive to Y1 receptors, highlighting that NPY has pathway-specific effects in hippocampal area CA1. This key finding may be the underlying cause of Y1 receptor agonism in CA1 causing anxiolytic effects. We also report that NPY overexpression causes a loss of Y1 receptor sensitivity in CA1, raising concerns that chronic NPY administration may also impair Y1 receptor function and thus have limited potential for the treatment of anxiety.

## References

Allen, Y.S., Adrian, T.E., Allen, J.M., Tatemoto, K., Crow, T.J., Bloom, S.R., Polak, J.M., 1983. Neuropeptide Y distribution in the rat brain. Science 221, 877–879.

Andriushchenko, N., Nebogina, K., Zorkina, Y., Abramova, O., Zubkov, E., Ochneva, A., Ushakova, V., Pavlov, K., Gurina, O., Chekhonin, V., Morozova, A., 2022. Antidepressant effect of neuropeptide Y in models of acute and chronic stress. Sci. Pharm. 90, 50. 10.3390/scipharm90030050.

Bale, R., Doshi, G., 2023. Cross talk about the role of Neuropeptide Y in CNS disorders and diseases. Neuropeptides 102, 102388. 10.1016/j.npep.2023.102388.

Bertocchi, I., Oberto, A., Longo, A., Mele, P., Sabetta, M., Bartolomucci, A., Palanza, P., Sprengel, R., Eva, C., 2011. Regulatory functions of limbic Y1 receptors in body weight and anxiety uncovered by conditional knockout and maternal care. Proc Natl Acad Sci USA 108, 19395–19400. 10.1073/pnas.1109468108.

Bjørnebekk, A., Mathé, A.A., Brené, S., 2010. The antidepressant effects of running and escitalopram are associated with levels of hippocampal NPY and Y1 receptor but not cell proliferation in a rat model of depression. Hippocampus 20, 820–828. 10.1002/hipo.20683.

Black, J.B., Premont, R.T., Daaka, Y., 2016. Feedback regulation of G protein-coupled receptor signaling by GRKs and arrestins. Semin. Cell Dev. Biol. 50, 95–104. 10.1016/j.semcdb.2015.12.015.

Cai, X., Kallarackal, A.J., Kvarta, M.D., Goluskin, S., Gaylor, K., Bailey, A.M., Lee, H.-K., Huganir, R.L., Thompson, S.M., 2013. Local potentiation of excitatory synapses by serotonin and its alteration in rodent models of depression. Nat. Neurosci. 16, 464–472. 10.1038/nn.3355.

Çalışkan, G., Stork, O., 2019. Hippocampal network oscillations at the interplay between innate anxiety and learned fear. Psychopharmacology (Berl) 236, 321–338. 10.1007/s00213-018-5109-z.

Cattaneo, S., Verlengia, G., Marino, P., Simonato, M., Bettegazzi, B., 2020. NPY and gene therapy for epilepsy: how, when,… and Y. Front. Mol. Neurosci. 13, 608001. 10.3389/fnmol.2020.608001.

Christiansen, S.H., Olesen, M.V., Gøtzsche, C.R., Woldbye, D.P.D., 2014. Anxiolytic-like effects after vector-mediated overexpression of neuropeptide Y in the amygdala and hippocampus of mice. Neuropeptides 48, 335–344. 10.1016/j.npep.2014.09.004.

Cohen, H., Liu, T., Kozlovsky, N., Kaplan, Z., Zohar, J., Mathé, A.A., 2012. The neuropeptide Y (NPY)-ergic system is associated with behavioral resilience to stress exposure in an animal model of post-traumatic stress disorder. Neuropsychopharmacology 37, 350–363. 10.1038/npp.2011.230.

Colmers, W.F., Klapstein, G.J., Fournier, A., St-Pierre, S., Treherne, K.A., 1991. Presynaptic inhibition by neuropeptide Y in rat hippocampal slice in vitro is mediated by a Y2 receptor. Br. J. Pharmacol. 102, 41–44. 10.1111/j.1476-5381.1991.tb12129.x.

Colmers, W.F., Lukowiak, K., Pittman, Q.J., 1988. Neuropeptide Y action in the rat hippocampal slice: site and mechanism of presynaptic inhibition. J. Neurosci. 8, 3827–3837.

Colmers, W.F., Lukowiak, K., Pittman, Q.J., 1987. Presynaptic action of neuropeptide Y in area CA1 of the rat hippocampal slice. J Physiol (Lond) 383, 285–299. 10.1113/jphysiol.1987.sp016409.

Colmers, W.F., Lukowiak, K., Pittman, Q.J., 1985. Neuropeptide Y reduces orthodromically evoked population spike in rat hippocampal CA1 by a possibly presynaptic mechanism. Brain Res. 346, 404–408. 10.1016/0006-8993(85)90880-7.

Cominski, T.P., Jiao, X., Catuzzi, J.E., Stewart, A.L., Pang, K.C.H., 2014. The role of the hippocampus in avoidance learning and anxiety vulnerability. Front. Behav. Neurosci. 8, 273. 10.3389/fnbeh.2014.00273.

Corder, K.M., Li, Q., Cortes, M.A., Bartley, A.F., Davis, T.R., Dobrunz, L.E., 2020. Overexpression of neuropeptide Y decreases responsiveness to neuropeptide Y. Neuropeptides 79, 101979. 10.1016/j.npep.2019.101979.

Dobrunz, L.E., Stevens, C.F., 1999. Response of hippocampal synapses to natural stimulation patterns. Neuron 22, 157–166. 10.1016/s0896-6273(00)80687-x.

Doods, H.N., Wieland, H.A., Engel, W., Eberlein, W., Willim, K.D., Entzeroth, M., Wienen, W., Rudolf, K., 1996. BIBP 3226, the first selective neuropeptide Y1 receptor antagonist: a review of its pharmacological properties. Regul. Pept. 65, 71–77. 10.1016/0167-0115(96)00074-2.

Dum, E., Fürtinger, S., Gasser, E., Bukovac, A., Drexel, M., Tasan, R., Sperk, G., 2017. Effective G-protein coupling of Y2 receptors along axonal fiber tracts and its relevance for epilepsy. Neuropeptides 61, 49–55. 10.1016/j.npep.2016.10.005.

El Bahh, B., Balosso, S., Hamilton, T., Herzog, H., Beck-Sickinger, A.G., Sperk, G., Gehlert, D.R., Vezzani, A., Colmers, W.F., 2005. The anti-epileptic actions of neuropeptide Y in the hippocampus are mediated by Y and not Y receptors. Eur. J. Neurosci. 22, 1417–1430. 10.1111/j.1460-9568.2005.04338.x.

El Bahh, B., Cao, J.Q., Beck-Sickinger, A.G., Colmers, W.F., 2002. Blockade of neuropeptide Y(2) receptors and suppression of NPY’s anti-epileptic actions in the rat hippocampal slice by BIIE0246. Br. J. Pharmacol. 136, 502–509. 10.1038/sj.bjp.0704751.

Engin, E., Smith, K.S., Gao, Y., Nagy, D., Foster, R.A., Tsvetkov, E., Keist, R., Crestani, F., Fritschy, J.-M., Bolshakov, V.Y., Hajos, M., Heldt, S.A., Rudolph, U., 2016. Modulation of anxiety and fear via distinct intrahippocampal circuits. eLife 5, e14120. 10.7554/eLife.14120.

Engin, E., Treit, D., 2007. The role of hippocampus in anxiety: intracerebral infusion studies. Behav. Pharmacol. 18, 365–374. 10.1097/FBP.0b013e3282de7929.

Enman, N.M., Sabban, E.L., McGonigle, P., Van Bockstaele, E.J., 2015. Targeting the Neuropeptide Y System in Stress-related Psychiatric Disorders. Neurobiol. Stress 1, 33–43. 10.1016/j.ynstr.2014.09.007.

Fekete, C., Sarkar, S., Rand, W.M., Harney, J.W., Emerson, C.H., Bianco, A.C., Beck-Sickinger, A., Lechan, R.M., 2002. Neuropeptide Y1 and Y5 receptors mediate the effects of neuropeptide Y on the hypothalamic-pituitary-thyroid axis. Endocrinology 143, 4513–4519. 10.1210/en.2002-220574.

Furtinger, S., Pirker, S., Czech, T., Baumgartner, C., Ransmayr, G., Sperk, G., 2001. Plasticity of Y1 and Y2 receptors and neuropeptide Y fibers in patients with temporal lobe epilepsy. J. Neurosci. 21, 5804–5812.

Gehlert, D.R., Schober, D.A., Morin, M., Berglund, M.M., 2007. Co-expression of neuropeptide Y Y1 and Y5 receptors results in heterodimerization and altered functional properties. Biochem. Pharmacol. 74, 1652–1664. 10.1016/j.bcp.2007.08.017.

Gicquiaux, H., Lecat, S., Gaire, M., Dieterlen, A., Mély, Y., Takeda, K., Bucher, B., Galzi, J.-L., 2002. Rapid internalization and recycling of the human neuropeptide Y Y(1) receptor. J. Biol. Chem. 277, 6645–6655. 10.1074/jbc.M107224200.

Giesbrecht, C.J., Mackay, J.P., Silveira, H.B., Urban, J.H., Colmers, W.F., 2010. Countervailing modulation of Ih by neuropeptide Y and corticotrophin-releasing factor in basolateral amygdala as a possible mechanism for their effects on stress-related behaviors. J. Neurosci. 30, 16970–16982. 10.1523/JNEUROSCI.2306-10.2010.

Goosens, K.A., 2011. Hippocampal regulation of aversive memories. Curr. Opin. Neurobiol. 21, 460–466. 10.1016/j.conb.2011.04.003.

Greber, S., Schwarzer, C., Sperk, G., 1994. Neuropeptide Y inhibits potassium-stimulated glutamate release through Y2 receptors in rat hippocampal slices in vitro. Br. J. Pharmacol. 113, 737–740. 10.1111/j.1476-5381.1994.tb17055.x.

Guilloux, J.P., Douillard-Guilloux, G., Kota, R., Wang, X., Gardier, A.M., Martinowich, K., Tseng, G.C., Lewis, D.A., Sibille, E., 2012. Molecular evidence for BDNF- and GABA-related dysfunctions in the amygdala of female subjects with major depression. Mol. Psychiatry 17, 1130–1142. 10.1038/mp.2011.113.

Hamilton, T.J., Xapelli, S., Michaelson, S.D., Larkum, M.E., Colmers, W.F., 2013. Modulation of distal calcium electrogenesis by neuropeptide Y₁ receptors inhibits neocortical long-term depression. J. Neurosci. 33, 11184–11193. 10.1523/JNEUROSCI.5595-12.2013.

Hashimoto, H., Onishi, H., Koide, S., Kai, T., Yamagami, S., 1996. Plasma neuropeptide Y in patients with major depressive disorder. Neurosci. Lett. 216, 57–60. 10.1016/0304-3940(96)13008-1.

Heilig, M., Söderpalm, B., Engel, J.A., Widerlöv, E., 1989. Centrally administered neuropeptide Y (NPY) produces anxiolytic-like effects in animal anxiety models. Psychopharmacology (Berl) 98, 524–529. 10.1007/BF00441953.

Heilig, M., 2004. The NPY system in stress, anxiety and depression. Neuropeptides 38, 213–224. 10.1016/j.npep.2004.05.002.

Hill, J.W., Urban, J.H., Xu, M., Levine, J.E., 2004. Estrogen Induces Neuropeptide Y (NPY) Y1 receptor gene expression and responsiveness to NPY in gonadotrope-enriched pituitary cell cultures. Endocrinology 145, 2283–2290. 10.1210/en.2003-1368.

Hirsch, S.J., Regmi, N.L., Birnbaum, S.G., Greene, R.W., 2015. CA1-specific deletion of NMDA receptors induces abnormal renewal of a learned fear response. Hippocampus 25, 1374–1379. 10.1002/hipo.22457.

Holliday, N.D., Lam, C.-W., Tough, I.R., Cox, H.M., 2005. Role of the C terminus in neuropeptide Y Y1 receptor desensitization and internalization. Mol. Pharmacol. 67, 655–664. 10.1124/mol.104.006114.

Hörmer, B.A., Verma, D., Gasser, E., Wieselthaler-Hölzl, A., Herzog, H., Tasan, R.O., 2018. Hippocampal NPY Y2 receptors modulate memory depending on emotional valence and time. Neuropharmacology 143, 20–28. 10.1016/j.neuropharm.2018.09.018.

Husum, H., Mathé, A.A., 2002. Early life stress changes concentrations of neuropeptide Y and corticotropin-releasing hormone in adult rat brain. Lithium treatment modifies these changes. Neuropsychopharmacology 27, 756–764. 10.1016/S0893-133X(02)00363-9.

Isomura, Y., Sirota, A., Ozen, S., Montgomery, S., Mizuseki, K., Henze, D.A., Buzsáki, G., 2006. Integration and segregation of activity in entorhinal-hippocampal subregions by neocortical slow oscillations. Neuron 52, 871–882. 10.1016/j.neuron.2006.10.023.

Jiménez-Vasquez, P.A., Diaz-Cabiale, Z., Caberlotto, L., Bellido, I., Overstreet, D., Fuxe, K., Mathé, A.A., 2007. Electroconvulsive stimuli selectively affect behavior and neuropeptide Y (NPY) and NPY Y(1) receptor gene expressions in hippocampus and hypothalamus of Flinders Sensitive Line rat model of depression. Eur. Neuropsychopharmacol. 17, 298–308. 10.1016/j.euroneuro.2006.06.011.

Jiménez-Vasquez, P.A., Overstreet, D.H., Mathé, A.A., 2000. Neuropeptide Y in male and female brains of Flinders Sensitive Line, a rat model of depression. Effects of electroconvulsive stimuli. J. Psychiatr. Res. 34, 405–412. 10.1016/S0022-3956(00)00036-4.

Kallarackal, A.J., Kvarta, M.D., Cammarata, E., Jaberi, L., Cai, X., Bailey, A.M., Thompson, S.M., 2013. Chronic stress induces a selective decrease in AMPA receptor-mediated synaptic excitation at hippocampal temporoammonic-CA1 synapses. J. Neurosci. 33, 15669–15674. 10.1523/JNEUROSCI.2588-13.2013.

Kalra, S.P., Dube, M.G., Fournier, A., Kalra, P.S., 1991. Structure-function analysis of stimulation of food intake by neuropeptide Y: effects of receptor agonists. Physiol. Behav. 50, 5–9. 10.1016/0031-9384(91)90490-f.

Karlsson, R.-M., Holmes, A., Heilig, M., Crawley, J.N., 2005. Anxiolytic-like actions of centrally-administered neuropeptide Y, but not galanin, in C57BL/6J mice. Pharmacol. Biochem. Behav. 80, 427–436. 10.1016/j.pbb.2004.12.009.

Karl, T., Burne, T.H.J., Herzog, H., 2006. Effect of Y1 receptor deficiency on motor activity, exploration, and anxiety. Behav. Brain Res. 167, 87–93. 10.1016/j.bbr.2005.08.019.

Kask, A., Vasar, E., Heidmets, L.T., Allikmets, L., Wikberg, J.E., 2001. Neuropeptide Y Y(5) receptor antagonist CGP71683A: the effects on food intake and anxiety-related behavior in the rat. Eur. J. Pharmacol. 414, 215–224. 10.1016/s0014-2999(01)00768-3.

Kautz, M., Charney, D.S., Murrough, J.W., 2017. Neuropeptide Y, resilience, and PTSD therapeutics. Neurosci. Lett. 649, 164–169. 10.1016/j.neulet.2016.11.061.

Kempf, N., Didier, P., Postupalenko, V., Bucher, B., Mély, Y., 2015. Internalization mechanism of neuropeptide Y bound to its Y1 receptor investigated by high resolution microscopy. Methods Appl. Fluoresc. 3, 025004. 10.1088/2050-6120/3/2/025004.

Klapstein, G.J., Colmers, W.F., 1997. Neuropeptide Y suppresses epileptiform activity in rat hippocampus in vitro. J. Neurophysiol. 78, 1651–1661. 10.1152/jn.1997.78.3.1651.

Kopp, J., Xu, Z.Q., Zhang, X., Pedrazzini, T., Herzog, H., Kresse, A., Wong, H., Walsh, J.H., Hökfelt, T., 2002. Expression of the neuropeptide Y Y1 receptor in the CNS of rat and of wild-type and Y1 receptor knock-out mice. Focus on immunohistochemical localization. Neuroscience 111, 443–532. 10.1016/s0306-4522(01)00463-8.

Kupcova, I., Danisovic, L., Grgac, I., Harsanyi, S., 2022. Anxiety and depression: what do we know of neuropeptides? Behav Sci (Basel) 12. 10.3390/bs12080262.

Kvarta, M.D., Bradbrook, K.E., Dantrassy, H.M., Bailey, A.M., Thompson, S.M., 2015. Corticosterone mediates the synaptic and behavioral effects of chronic stress at rat hippocampal temporoammonic synapses. J. Neurophysiol. 114, 1713–1724. 10.1152/jn.00359.2015.

Ledoux, V.A., Smejkalova, T., May, R.M., Cooke, B.M., Woolley, C.S., 2009. Estradiol facilitates the release of neuropeptide Y to suppress hippocampus-dependent seizures. J. Neurosci. 29, 1457–1468. 10.1523/JNEUROSCI.4688-08.2009.

Ledri, M., Sørensen, A.T., Madsen, M.G., Christiansen, S.H., Ledri, L.N., Cifra, A., Bengzon, J., Lindberg, E., Pinborg, L.H., Jespersen, B., Gøtzsche, C.R., Woldbye, D.P.D., Andersson, M., Kokaia, M., 2015. Differential effect of neuropeptides on excitatory synaptic transmission in human epileptic hippocampus. J. Neurosci. 35, 9622–9631. 10.1523/JNEUROSCI.3973-14.2015.

Li, Q., Bartley, A.F., Dobrunz, L.E., 2017. Endogenously Released Neuropeptide Y Suppresses Hippocampal Short-Term Facilitation and Is Impaired by Stress-Induced Anxiety. J. Neurosci. 37, 23–37. 10.1523/JNEUROSCI.2599-16.2016.

Longo, A., Mele, P., Bertocchi, I., Oberto, A., Bachmann, A., Bartolomucci, A., Palanza, P., Sprengel, R., Eva, C., 2014. Conditional inactivation of neuropeptide Y Y1 receptors unravels the role of Y1 and Y5 receptors coexpressing neurons in anxiety. Biol. Psychiatry 76, 840–849. 10.1016/j.biopsych.2014.01.009.

Longo, A., Oberto, A., Mele, P., Mattiello, L., Pisu, M.G., Palanza, P., Serra, M., Eva, C., 2015. NPY-Y1 coexpressed with NPY-Y5 receptors modulate anxiety but not mild social stress response in mice. Genes Brain Behav. 14, 534–542. 10.1111/gbb.12232.

Lörincz, A., Notomi, T., Tamás, G., Shigemoto, R., Nusser, Z., 2002. Polarized and compartment-dependent distribution of HCN1 in pyramidal cell dendrites. Nat. Neurosci. 5, 1185–1193. 10.1038/nn962.

Lovett-Barron, M., Kaifosh, P., Kheirbek, M.A., Danielson, N., Zaremba, J.D., Reardon, T.R., Turi, G.F., Hen, R., Zemelman, B.V., Losonczy, A., 2014. Dendritic inhibition in the hippocampus supports fear learning. Science 343, 857–863. 10.1126/science.1247485.

Marvizon, J.C., Chen, W., Fu, W., Taylor, B.K., 2019. Neuropeptide Y release in the rat spinal cord measured with Y1 receptor internalization is increased after nerve injury. Neuropharmacology 158, 107732. 10.1016/j.neuropharm.2019.107732.

Ma, T.-L., Zhou, Y., Zhang, C.-Y., Gao, Z.-A., Duan, J.-X., 2021. The role and mechanism of β-arrestin2 in signal transduction. Life Sci. 275, 119364. 10.1016/j.lfs.2021.119364.

Mathé, A.A., Michaneck, M., Berg, E., Charney, D.S., Murrough, J.W., 2020. A randomized controlled trial of intranasal neuropeptide Y in patients with major depressive disorder. Int. J. Neuropsychopharmacol. 23, 783–790. 10.1093/ijnp/pyaa054.

McQuiston, A.R., Colmers, W.F., 1996. Neuropeptide Y2 receptors inhibit the frequency of spontaneous but not miniature EPSCs in CA3 pyramidal cells of rat hippocampus. J. Neurophysiol. 76, 3159–3168. 10.1152/jn.1996.76.5.3159.

McQuiston, A.R., Petrozzino, J.J., Connor, J.A., Colmers, W.F., 1996. Neuropeptide Y1 receptors inhibit N-type calcium currents and reduce transient calcium increases in rat dentate granule cells. J. Neurosci. 16, 1422–1429. 10.1523/JNEUROSCI.16-04-01422.1996.

Méndez-Couz, M., Manahan-Vaughan, D., Silva, A.P., González-Pardo, H., Arias, J.L., Conejo, N.M., 2021. Metaplastic contribution of neuropeptide Y receptors to spatial memory acquisition. Behav. Brain Res. 396, 112864. 10.1016/j.bbr.2020.112864.

Molosh, A.I., Sajdyk, T.J., Truitt, W.A., Zhu, W., Oxford, G.S., Shekhar, A., 2013. NPY Y1 receptors differentially modulate GABAA and NMDA receptors via divergent signal-transduction pathways to reduce excitability of amygdala neurons. Neuropsychopharmacology 38, 1352–1364. 10.1038/npp.2013.33.

Morales-Medina, J.C., Dumont, Y., Quirion, R., 2010. A possible role of neuropeptide Y in depression and stress. Brain Res. 1314, 194–205. 10.1016/j.brainres.2009.09.077.

Nahvi, R.J., Sabban, E.L., 2020. Sex differences in the neuropeptide Y system and implications for stress related disorders. Biomolecules 10. 10.3390/biom10091248.

Nelson, T.S., Taylor, B.K., 2021. Targeting spinal neuropeptide Y1 receptor-expressing interneurons to alleviate chronic pain and itch. Prog. Neurobiol. 196, 101894. 10.1016/j.pneurobio.2020.101894.

Olesen, M.V., Christiansen, S.H., Gøtzsche, C.R., Holst, B., Kokaia, M., Woldbye, D.P.D., 2012. Y5 neuropeptide Y receptor overexpression in mice neither affects anxiety- and depression-like behaviours nor seizures but confers moderate hyperactivity. Neuropeptides 46, 71–79. 10.1016/j.npep.2012.01.002.

Parker, R.M., Herzog, H., 1999. Regional distribution of Y-receptor subtype mRNAs in rat brain. Eur. J. Neurosci. 11, 1431–1448. 10.1046/j.1460-9568.1999.00553.x.

Parker, R.M., Herzog, H., 1998. Comparison of Y-receptor subtype expression in the rat hippocampus. Regul. Pept. 75–76, 109–115. 10.1016/s0167-0115(98)00059-7.

Pickel, V.M., Chan, J., Veznedaroglu, E., Milner, T.A., 1995. Neuropeptide Y and dynorphin-immunoreactive large dense-core vesicles are strategically localized for presynaptic modulation in the hippocampal formation and substantia nigra. Synapse 19, 160–169. 10.1002/syn.890190303.

Pleil, K.E., Rinker, J.A., Lowery-Gionta, E.G., Mazzone, C.M., McCall, N.M., Kendra, A.M., Olson, D.P., Lowell, B.B., Grant, K.A., Thiele, T.E., Kash, T.L., 2015. NPY signaling inhibits extended amygdala CRF neurons to suppress binge alcohol drinking. Nat. Neurosci. 18, 545–552. 10.1038/nn.3972.

Pronchuk, N., Beck-Sickinger, A.G., Colmers, W.F., 2002. Multiple NPY receptors Inhibit GABA(A) synaptic responses of rat medial parvocellular effector neurons in the hypothalamic paraventricular nucleus. Endocrinology 143, 535–543. 10.1210/endo.143.2.8655.

Rasmusson, A.M., Pineles, S.L., 2018. Neurotransmitter, peptide, and steroid hormone abnormalities in PTSD: biological endophenotypes relevant to treatment. Curr. Psychiatry Rep. 20, 52. 10.1007/s11920-018-0908-9.

Redrobe, J.P., Dumont, Y., Fournier, A., Quirion, R., 2002a. The neuropeptide Y (NPY) Y1 receptor subtype mediates NPY-induced antidepressant-like activity in the mouse forced swimming test. Neuropsychopharmacology 26, 615–624. 10.1016/S0893-133X(01)00403-1.

Redrobe, J.P., Dumont, Y., Quirion, R., 2002b. Neuropeptide Y (NPY) and depression: from animal studies to the human condition. Life Sci. 71, 2921–2937. 10.1016/S0024-3205(02)02159-8.

Reichmann, F., Holzer, P., 2016. Neuropeptide Y: A stressful review. Neuropeptides 55, 99–109. 10.1016/j.npep.2015.09.008.

Sah, R., Ekhator, N.N., Jefferson-Wilson, L., Horn, P.S., Geracioti, T.D., 2014. Cerebrospinal fluid neuropeptide Y in combat veterans with and without posttraumatic stress disorder. Psychoneuroendocrinology 40, 277–283. 10.1016/j.psyneuen.2013.10.017.

Sajdyk, T.J., Johnson, P.L., Leitermann, R.J., Fitz, S.D., Dietrich, A., Morin, M., Gehlert, D.R., Urban, J.H., Shekhar, A., 2008. Neuropeptide Y in the amygdala induces long-term resilience to stress-induced reductions in social responses but not hypothalamic-adrenal-pituitary axis activity or hyperthermia. J. Neurosci. 28, 893–903. 10.1523/JNEUROSCI.0659-07.2008.

Sayed, S., Van Dam, N.T., Horn, S.R., Kautz, M.M., Parides, M., Costi, S., Collins, K.A., Iacoviello, B., Iosifescu, D.V., Mathé, A.A., Southwick, S.M., Feder, A., Charney, D.S., Murrough, J.W., 2018. A Randomized Dose-Ranging Study of Neuropeptide Y in Patients with Posttraumatic Stress Disorder. Int. J. Neuropsychopharmacol. 21, 3–11. 10.1093/ijnp/pyx109.

Schmeltzer, S.N., Herman, J.P., Sah, R., 2016. Neuropeptide Y (NPY) and posttraumatic stress disorder (PTSD): A translational update. Exp. Neurol. 284, 196–210. 10.1016/j.expneurol.2016.06.020.

Silva, A.P., Carvalho, A.P., Carvalho, C.M., Malva, J.O., 2001. Modulation of intracellular calcium changes and glutamate release by neuropeptide Y1 and Y2 receptors in the rat hippocampus: differential effects in CA1, CA3 and dentate gyrus. J. Neurochem. 79, 286–296. 10.1046/j.1471-4159.2001.00560.x.

Silveira, M.A., Drotos, A.C., Pirrone, T.M., Versalle, T.S., Bock, A., Roberts, M.T., 2023. Neuropeptide Y signaling regulates recurrent excitation in the auditory midbrain. J. Neurosci. 43, 7626–7641. 10.1523/JNEUROSCI.0900-23.2023.

Silveira Villarroel, H., Bompolaki, M., Mackay, J.P., Miranda Tapia, A.P., Michaelson, S.D., Leitermann, R.J., Marr, R.A., Urban, J.H., Colmers, W.F., 2018. NPY induces stress resilience via downregulation of ih in principal neurons of rat basolateral amygdala. J. Neurosci. 38, 4505–4520. 10.1523/JNEUROSCI.3528-17.2018.

Sippy, T., Cruz-Martín, A., Jeromin, A., Schweizer, F.E., 2003. Acute changes in short-term plasticity at synapses with elevated levels of neuronal calcium sensor-1. Nat. Neurosci. 6, 1031–1038. 10.1038/nn1117.

Smiałowska, M., Wierońska, J.M., Domin, H., Zieba, B., 2007. The effect of intrahippocampal injection of group II and III metobotropic glutamate receptor agonists on anxiety; the role of neuropeptide Y. Neuropsychopharmacology 32, 1242–1250. 10.1038/sj.npp.1301258.

Smith, S.J., Sümbül, U., Graybuck, L.T., Collman, F., Seshamani, S., Gala, R., Gliko, O., Elabbady, L., Miller, J.A., Bakken, T.E., Rossier, J., Yao, Z., Lein, E., Zeng, H., Tasic, B., Hawrylycz, M., 2019. Single-cell transcriptomic evidence for dense intracortical neuropeptide networks. eLife 8. 10.7554/eLife.47889.

Sommer, W.H., Lidström, J., Sun, H., Passer, D., Eskay, R., Parker, S.C.J., Witt, S.H., Zimmermann, U.S., Nieratschker, V., Rietschel, M., Margulies, E.H., Palkovits, M., Laucht, M., Heilig, M., 2010. Human NPY promoter variation rs16147:T>C as a moderator of prefrontal NPY gene expression and negative affect. Hum. Mutat. 31, E1594–608. 10.1002/humu.21299.

Sperk, G., Hamilton, T., Colmers, W.F., 2007. Neuropeptide Y in the dentate gyrus. Prog. Brain Res. 163, 285–297. 10.1016/S0079-6123(07)63017-9.

Ste Marie, L., Luquet, S., Cole, T.B., Palmiter, R.D., 2005. Modulation of neuropeptide Y expression in adult mice does not affect feeding. Proc Natl Acad Sci USA 102, 18632–18637. 10.1073/pnas.0509240102.

Sun, H.Y., Li, Q., Bartley, A.F., Dobrunz, L.E., 2018. Target-cell-specific Short-term Plasticity Reduces the Excitatory Drive onto CA1 Interneurons Relative to Pyramidal Cells During Physiologically-derived Spike Trains. Neuroscience 388, 430–447. 10.1016/j.neuroscience.2018.07.051.

Tural, U., Iosifescu, D.V., 2020. Neuropeptide Y in PTSD, MDD, and chronic stress: A systematic review and meta-analysis. J. Neurosci. Res. 98, 950–963. 10.1002/jnr.24589.

van den Pol, A.N., 2012. Neuropeptide transmission in brain circuits. Neuron 76, 98–115. 10.1016/j.neuron.2012.09.014.

Walters, B.J., Hallengren, J.J., Theile, C.S., Ploegh, H.L., Wilson, S.M., Dobrunz, L.E., 2014. A catalytic independent function of the deubiquitinating enzyme USP14 regulates hippocampal synaptic short-term plasticity and vesicle number. J Physiol (Lond) 592, 571–586. 10.1113/jphysiol.2013.266015.

Widerlöv, E., Lindström, L.H., Wahlestedt, C., Ekman, R., 1988. Neuropeptide Y and peptide YY as possible cerebrospinal fluid markers for major depression and schizophrenia, respectively. J. Psychiatr. Res. 22, 69–79. 10.1016/0022-3956(88)90030-1.

Wolak, M.L., DeJoseph, M.R., Cator, A.D., Mokashi, A.S., Brownfield, M.S., Urban, J.H., 2003. Comparative distribution of neuropeptide Y Y1 and Y5 receptors in the rat brain by using immunohistochemistry. J. Comp. Neurol. 464, 285–311. 10.1002/cne.10823.

Wu, G., Feder, A., Wegener, G., Bailey, C., Saxena, S., Charney, D., Mathé, A.A., 2011. Central functions of neuropeptide Y in mood and anxiety disorders. Expert Opin. Ther. Targets 15, 1317–1331. 10.1517/14728222.2011.628314.

Zhou, Z., Zhu, G., Hariri, A.R., Enoch, M.-A., Scott, D., Sinha, R., Virkkunen, M., Mash, D.C., Lipsky, R.H., Hu, X.-Z., Hodgkinson, C.A., Xu, K., Buzas, B., Yuan, Q., Shen, P.-H., Ferrell, R.E., Manuck, S.B., Brown, S.M., Hauger, R.L., Stohler, C.S., Zubieta, J.-K., Goldman, D., 2008. Genetic variation in human NPY expression affects stress response and emotion. Nature 452, 997–1001. 10.1038/nature06858.

Zucker, R.S., Delaney, K.R., Mulkey, R., Tank, D.W., 1991. Presynaptic calcium in transmitter release and posttetanic potentiation. Ann. N. Y. Acad. Sci. 635, 191–207. 10.1111/j.1749-6632.1991.tb36492.x.

